# Time to run: Late rather than early exercise training in mice remodels the gut microbiome and reduces atherosclerosis development

**DOI:** 10.1101/2022.11.28.518180

**Authors:** Milena Schönke, Zhixiong Ying, Artemiy Kovynev, Wietse In het Panhuis, Anne Binnendijk, Sabine van der Poel, Amanda C.M. Pronk, Trea C.M Streefland, Menno Hoekstra, Sander Kooijman, Patrick C.N. Rensen

## Abstract

The metabolic and inflammatory processes that are implicated in the development of cardiovascular diseases are under control of the biological clock. While skeletal muscle function exhibits circadian rhythms, it is unclear to what extent the beneficial health effects of exercise are restricted to unique time windows. We aimed to study whether the timing of exercise training differentially modulates the development of atherosclerosis and elucidate underlying mechanisms. We endurance-trained atherosclerosis-prone female APOE*3-Leiden.CETP mice fed a Western-type diet, a well-established human-like model for cardiometabolic diseases, for one hour five times a week for four weeks either in their early or in their late active phase on a treadmill. We monitored metabolic parameters, the development of atherosclerotic lesions in the aortic root and assessed the composition of the gut microbiota. Late, but not early, exercise training reduced fat mass by 19% and the size of early-stage atherosclerotic lesions by as much as 29% compared to sedentary animals. No correlation between cholesterol exposure and lesion size was evident, as no differences in plasma lipid levels were observed, but circulating levels of the pro-inflammatory markers ICAM-1 and VCAM-1 were reduced with late exercise. Strikingly, we observed a time-of-day-dependent effect of exercise training on the composition of the gut microbiota as only late training increased the abundance of gut bacteria producing short-chain fatty acids with proposed anti-inflammatory properties. Together, these findings indicate that timing is a critical factor to the beneficial anti-atherosclerotic effects of exercise with a great potential to further optimize training recommendations for patients.

## 1. Introduction

Cardiovascular diseases (CVD) remain the leading cause of death worldwide, taking nearly 18 million lives each year. A cornerstone of the prevention and management of the main underlying cause of CVD, atherosclerosis, is an active lifestyle with regular exercise. In patients enrolled in exercise training programs a significant improvement of cardiorespiratory fitness and cardiometabolic risk factors such as reduced plasma lipid levels and fasting insulin can be observed [1, 2].

Many of the molecular processes dysregulated in CVD development, including glucose and lipid metabolism, whole-body inflammation and the interaction with the gut microbiome, are under the control of the circadian clock [3, 4]. While it is well established that chronic circadian disruption, for instance seen in shift workers, is detrimental to cardiometabolic health [5], it is currently not entirely understood at what time of day exercise aligns best with the intrinsic circadian control and what the “ideal” training time is [6]. Exercise performance has long been known to peak in the evening which coincides with the maximum mitochondrial capacity of skeletal muscle [7, 8], but it remains unclear whether this is also the best time for exercise training with regard to the prevention and treatment of common cardiometabolic diseases. We previously demonstrated in mice that exercise has a unique time-dependent molecular fingerprint in skeletal muscle and other tissues like liver, suggesting that the right timing of exercise may be crucial to maximize metabolic health benefits [9, 10]. Identifying the “best” time to exercise would not only enhance the efficacy of exercise programs for CVD patients but potentially also reduce the need for drug treatment and the number of individuals becoming patients in the first place.

A randomized crossover trial with men with type 2 diabetes found that high-intensity interval cycling training in the afternoon was more efficacious in lowering glycemia than training in the morning which even had a deleterious effect on blood glucose regulation [11]. Similarly, only evening exercise improved fasting blood glucose, insulin, cholesterol, triglyceride (TG) and low-density lipoprotein (LDL)-cholesterol concentrations in the blood of men with obesity following a short-time high fat diet intervention as well as ambulatory blood pressure in men with treated hypertension [12, 13].

Studying atherosclerosis development in clinical intervention studies is difficult as the disease develops over years or decades and regresses only slowly. Several mouse models are commonly used in the study of atherosclerosis and it has been demonstrated that exercise training reduces atherosclerosis in both LDL receptor knockout as well ApoE knockout animals [14-16]. These mice develop hypercholesterolemia and subsequently atherosclerotic lesions in the walls of arteries as a consequence of an interrupted ApoE-LDL receptor pathway responsible for the clearance of atherogenic lipoprotein remnants via the liver. However, exercise may influence this pathway and modulate whole-body lipid and lipoprotein metabolism through the increased uptake of TG-derived fatty acids by active skeletal muscle. Accordingly, we chose APOE*3-Leiden.CETP mice that possess an intact ApoE-LDL receptor pathway and a humanized lipoprotein profile to study whether the timing of aerobic exercise training matters for its anti-atherosclerotic effect [17]. The aim of this study was to identify the best time to train for the mitigation of CVD.

## 2. Methods

### 2.1 Animal study

Atherosclerosis-prone female APOE*3-Leiden.CETP mice were obtained as previously described [18]. At 8-12 weeks of age, mice were group-housed in light-tight cabinets at 21°C under 12/12-hour light-dark conditions with cage enrichment. The cabinets were illuminated with white fluorescent light (intensity: 200-250 lux). All training bouts and experiments took place under dim red light illumination during the dark (active) phase of the mice. Mice were fed a Western-type diet (containing 16% fat and 0.4% cholesterol; Ssniff, Soest, Germany) *ad libitum*. After a dietary habituation period of 10 days, animals were block-randomized into three groups based on body weight, fat mass, lean mass (assessed by EchoMRI 100-Analyzer; EchoMRI, Houston, Texas), plasma TG, and total plasma cholesterol (TC). Group sizes were decided based on effect size on atherosclerosis development observed in previous studies with this animal model. Animals without increased plasma lipids following the habituation period were excluded. Early dark phase running (“Early runners”, E-RUN, n=16) took place one hour after lights off at *Zeitgeber* time (ZT) 13-14, late dark phase running (“Late runners”, L-RUN, n=16) one hour before lights on at ZT 22-23 and sedentary mice (SED, n=16) did not train. The sedentary control group was split up, half (n=8) was housed in the same light cabinet as E-RUN and half (n=8) in the same light cabinet as L-RUN. They are shown together as SED in Fig. 1-3B and separated as E-SED and L-SED in Fig. 3C-5. Body weight of all mice was assessed weekly and body composition, unfasted TG and TC after four weeks at the end of the study. The animals were killed via CO_2_ inhalation 24 hours after their last training bout and perfused with ice-cold PBS for 5 minutes before tissues were collected for further analyses. The SED animals were killed at the same ZT times as their respective exercising counterparts. In a follow-up study with n=8 mice per group and the same training schedule, unfasted plasma glucose was measured directly after training and again after 4, 9 and 13 hours during the third week of training. Mice were single-housed in metabolic cages (Promethion, Sables Systems International, Las Vegas, Nevada) during the fourth week of training for 6 days including 2 days of acclimatization and plasma clearance and organ uptake of radiolabeled TG packaged within very low-density lipoprotein (VLDL)-like particles were assessed immediately after the last training bout after six weeks of training. All other experimental data shown here refers to the first cohort. All animal experiments were performed in accordance with the Institute for Laboratory Animal Research Guide for the Care and Use of Laboratory Animals and were approved by the National Committee for Animal experiments by the Ethics Committee on Animal Care and Experimentation of the Leiden University Medical Center.

**Figure 1.**
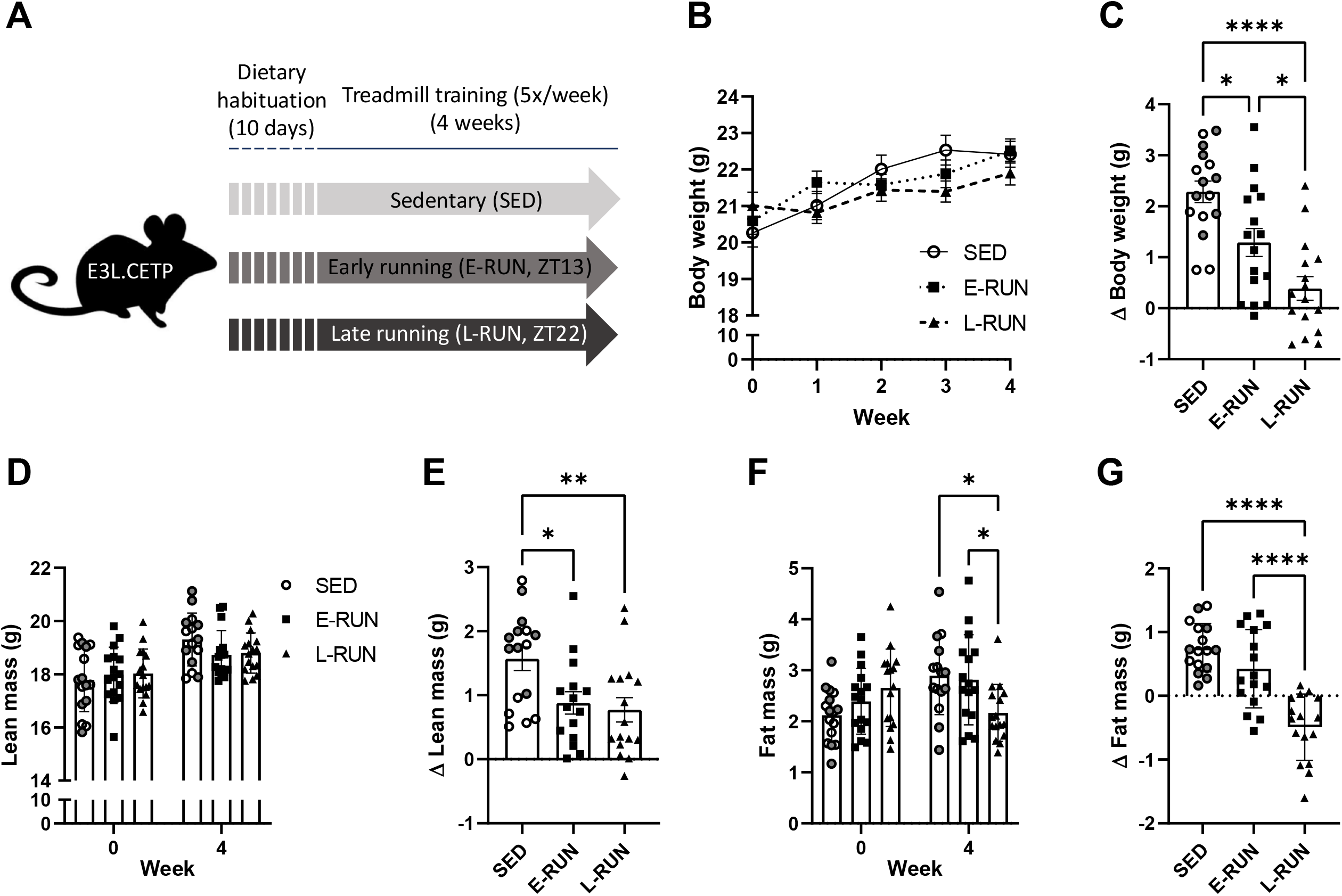
Only late exercise training promotes a loss of fat mass. Atherosclerosis-prone APOE*3-Leiden.CETP mice on a cholesterol-rich Western-type diet were treadmill trained five times per week either in the early or late dark phase for four weeks (**A**). Bodyweight (**B**), relative changes to body weight (**C**), lean mass (**D**), relative changes to lean mass (**E**), fat mass (**F**) and relative changes to fat mass (**G**) were monitored. Data are presented as means ± SEM (n= 15-16), *P<0.05, **P<0.01, ****P<0.0001 according to one-way ANOVA or mixed-effects analysis. E-SED data points are labeled white and L-SED data points dark grey within the SED group.

### 2.2 Exercise training

The mice were trained on a rat treadmill with five lanes (MazeEngineers, Skokie, Illinois), allowing 3-4 cage mates to run on one lane together which was observed to improve the motivation of the mice to run, when compared with solitary running. No electricity was used as stimulus; instead mice were gently nudged back on the belt with a brush when they stopped running. After three days of treadmill acclimatization with increasing speed and running duration the mice were trained five times per week for one hour (15 min warm-up at 6-15 m/min, 45 min at 15 m/min; in total 839 m per bout for each mouse) for a total of four weeks in the main atherosclerosis study and six weeks in the follow-up study. To account for handling stress and the unknown environment that the training mice experienced, all SED mice were moved in groups into empty cages without bedding standing close to the treadmill during the hour their exercising counterparts were trained.

### 2.3 Locomotor activity, food intake and energy expenditure measurements

Mice were individually housed in automated metabolic home cages (Promethion System, Sable Systems, Berlin, Germany) for 6 days including 2 days of acclimatization. O_2_ consumption, CO_2_ production, food intake and voluntary locomotor activity (by beam breaks) were continuously measured in 5-minute bins and energy expenditure, fat oxidation rate, carbohydrate oxidation rate and respiratory exchange ratio (RER) were calculated. The data from the last 4 days is shown both without normalization and with normalization to lean body mass.

### 2.4 Plasma lipid and glucose measurements

Plasma TG and TC were measured after the dietary habituation period directly before the start of the training and after four weeks using the Total Triglyceride and Total Cholesterol assay kits (both Roche Diagnostics, Almere, The Netherlands), respectively, as described previously [19]. High-density lipoprotein-cholesterol (HDL-C) was measured after four weeks of training. For this, ApoB-containing lipoproteins were precipitated as described elsewhere [19] and HDL-C was determined by measuring TC in the supernatant as described above. TC exposure was calculated by multiplying the average plasma TC concentrations (mM) throughout the study by the number of study weeks. Plasma glucose was measured using the Glucose liquiUV mono kit according to the handbook (Human, Wiesbaden, Germany).

### 2.5 TG-derived fatty acid uptake by organs

TG-rich VLDL-like particles (approx. 80 nm diameter) labeled with glycerol tri[^3^H]oleate were prepared as previously described [20]. In the follow-up study, after a baseline blood collection from the tail vein (t=0), mice were injected with VLDL-like particles intravenously via the tail vein (1 mg TG diluted in 200 μL saline per mouse) directly after the last running bout after six weeks of training to measure the uptake of TG-derived fatty acids by various tissues. Plasma was collected over time to determine glycerol tri[^3^H]oleate plasma decay. After 15 minutes, animals were killed by CO_2_ inhalation and perfused with ice-cold PBS for 5 minutes before tissues were collected for further analyses. Tissue samples for radioactivity measurement were prepared as described elsewhere [19].

### 2.6 Atherosclerosis quantification

Hearts were fixated in 4% paraformaldehyde for 24 hours and transferred to 70% ethanol before they were embedded in paraffin, cross-sectioned using a microtome (5 μm) and stained with hematoxylin-phloxine-saffron (HPS). Atherosclerotic lesion sizes of four cross sections throughout the aortic root with a distance of 50 μm, starting at open aortic valves, were quantified with imaging software while blinded (ImageJ version 1.52p; National Institutes of Health, Bethesda, Maryland). Atherosclerotic plaques were scored subjectively for lesion severity while blinded (Mild: type I-III lesions; Severe: type IV-V lesions), according to the guidelines of the American Heart Association adapted for mice [21]. A staining of macrophages was performed using a primary antibody against Mac3 (#550292, BD Biosciences, Franklin Lakes, New Jersey, [22]) 1:1000 and secondary ImmPRESS HRP Goat Anti-Rat IgG (Vector Laboratories, Burlingame, California) to assess the macrophage content of type III lesions.

### 2.7 Cytokine measurements

Interleukin (IL) 6, IL-1β, IL-10 and tumor necrosis factor (TNF) α were measured in plasma samples collected in the atherosclerosis cohort after four weeks of training using a customized multi-plex U-Plex Biomarker Group 1 (mouse) Kit (Meso Scale Discovery; Rockville, Maryland) according to the manufacturer’s protocol. Plasma ICAM-1 and VCAM-1 were measured using the Mouse ICAM-1/CD54 and VCAM-1/CD106 Quantikine ELISA kits according to the handbooks (R&D Systems, Minneapolis, USA).

### 2.8 Gene expression analysis

RNA was isolated from freshly frozen tissue (aortic arch and descending aorta; tibialis anterior and soleus muscle) using TRIzol RNA isolation reagent (Thermo Fisher, Waltham, Massachusetts). Following reverse transcription with M-MLV Reverse Transcriptase (Promega, Madison, Wisconsin), qRT-PCR was performed using SYBR Green (Promega). The primers used were (5’-3’) *Gpx1*: fw GGTTCGAGCCCAATTTTACA, rev CATTCCGCAGGAAGGTAAAG; *Sod1*: fw TACACAAGGCTGTACCAGTGC, rev ACATGCCTCTCTTCATCCGC; *Tnfa*: fw GCCTCTTCTCATTCCTGCTTG, rev CTGATGAGAGGGAGGCCATT; *Il1b*: fw GCAACTGTTCCTGAACTCAACT, rev ATCTTTTGGGGTCCGTCAACT; *Icam1*: fw TCCGCTGTGCTTTGAGAACT, rev TCCGGAAACGAATACACGGT; *Vcam1*: fw TGGAGGTCTACTCATTCCCTGA, rev GACAGGTCTCCCATGCACAA; *Olfr78*: fw GGCATTTGGGACTTGTGTGT, rev AGCACCGTAGATGATGGGATT; *Hif1a*: fw ACGAGAAGAAAAATAGGATGAGTTC, rev GTGGCAACTGATGAGCAAGC; *Cd163*: fw CTCAGGAAACCAATCCCAGA, rev CAAGAGCCCTCGTGGTAGAC and *Adgre1*: fw CTTTGGCTATGGGCTTCCAGTC, rev GCAAGGAGGACAGAGTTTATCGTG. Expression of genes of interest was normalized to expression of the housekeeping gene *Rplp0* (fw GGACCCGAGAAGACCTCCTT, rev GCACATCACTCAGAATTTCAATGG).

### 2.9 Gut microbiota sequencing

Total genomic DNA from cecum content (n=14-16 per group due to insufficient amount of cecum content) was isolated using the QIAamp DNA Stool Mini Kit from Qiagen (Venlo, The Netherlands). The hypervariable regions V3-V4 of the 16S rRNA coding gene were amplified using Illumina primers (fw: CCTACGGGNGGCWGCAG, rev: GACTACHVGGGTATCTAATCC) and sequencing (2 × 300 bp paired-end) was carried out on the MiSeq platform (Illumina, San Diego, California) with a coverage of 50k reads. The QIIME 2 2021.8 tool was used to process the data and statistical analyses were performed in R version 4.1.1 (both described in detail in the supplementary methods). On a subset of cecum content DNA samples (n=4-5 per group), metagenomic shotgun sequencing was performed on the HiSeq X Ten platform (Illumina). These data were processed using NGLess version 1.0 with the mOTU-tool (version 2.6) module and statistical analyses were performed in R version 4.1.1 (both described in detail in the supplementary methods).

### 2.10 Statistical analysis

All individual data points are shown or data are otherwise expressed as mean ± SEM. Statistical analyses were performed using GraphPad Prism 9.01 (GraphPad, La Jolla, California) and one-way or two-way ANOVAs followed by Tukey’s multiple comparisons test were used where appropriate. In case of missing values, mixed-effects analysis was performed. The data was tested for normality. Statistical outliers were removed after identification by Grubb’s test. Differences between groups were considered statistically significant if p < 0.05 (*), p < 0.01 (**), p < 0.001 (***) or p < 0.0001 (****).

## 3. Results

### 3.1 Only late exercise training causes a loss of body fat mass

APOE*3-Leiden.CETP mice were fed a cholesterol-rich atherosclerosis-inducing Western-type diet and treadmill trained for one hour on five days per week for four weeks either in the early (ZT 13) or late (ZT 22) dark phase (Fig. 1A) to mimic morning and evening exercise in humans. Both late runners (L-RUN) and early runners (E-RUN) gained significantly less body weight than sedentary animals (SED), and L-RUN gained less weight than E-RUN (Fig. 1B, C). This was due to both running groups gaining less lean mass than SED mice (Fig. 1D, E) and L-RUN in addition losing fat mass in comparison to both other groups over the four-week period (Fig. 1F, G; p=0.045 L-RUN week 0 vs week 4; SED +36.8 ± 4.7%, E-RUN +19.7 ± 7.8%, L-RUN -15.8 ± 4.2% over four weeks). As a marker of chronic stress exposure the weight of the adrenal glands was recorded and no differences between the groups were observed (Suppl. Fig. S1).

### 3.2 Only late exercise training reduces the development of atherosclerosis without ameliorating hyperlipidemia

Plasma TG, TC as well as HDL-C were not changed after four weeks of either early or late training in comparison to before the start of the training and to SED (Fig. 2A-C). Accordingly, all mice had a comparable cholesterol exposure throughout the study (Fig. 2D). Nonetheless, exercise training significantly reduced the size of atherosclerotic lesions in the aortic root of L-RUN, while the lesion burden of E-RUN was similar to that of SED (Fig. 2E, F, H). The lesion burden of SED housed with E-RUN and SED housed with L-RUN was not different (separated data not shown, p=0.97). Overall, the majority of atherosclerotic lesions observed were mild but L-RUN showed a tendency towards a greater relative amount of the mildest lesions (p=0.11, type I, early fatty streak in the intima of the vessel wall) and a significant reduction of type III lesions (extension of foam cells into media, lesion shows a fibrotic cap) as compared to E-RUN (Fig. 2G).

**Figure 2.**
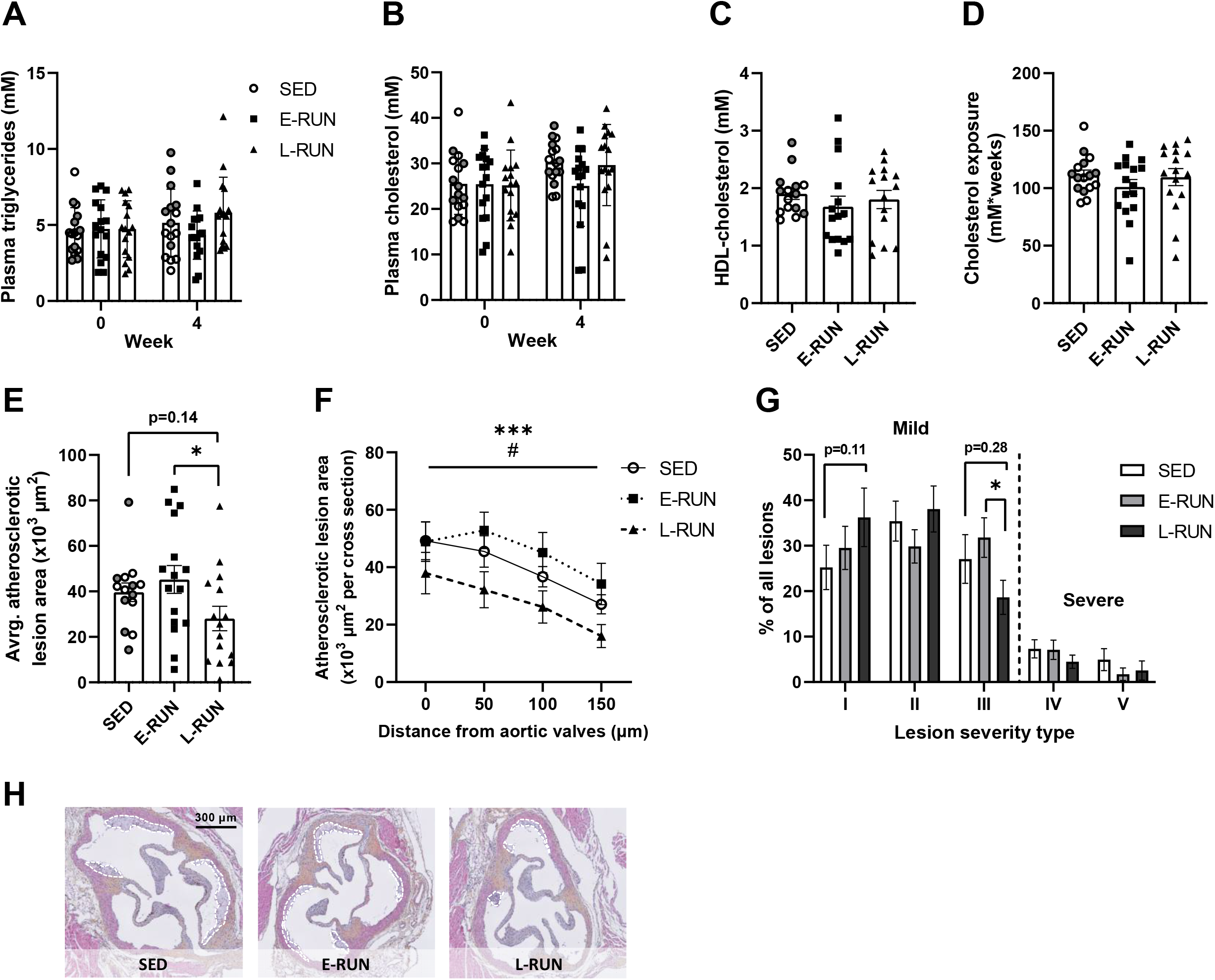
Only late exercise training reduces the development of atherosclerosis without ameliorating plasma lipid levels. At the beginning and the end of the training program, plasma triglyceride (**A**) and total cholesterol levels (**B**) were measured in blood collected before the first and 24 h after the last treadmill running bout. After four weeks of training, plasma HDL-cholesterol was measured (**C**) and the total cholesterol exposure over the four-week period was calculated (**D**). Average atherosclerotic lesion size in the aortic root (**E**) was assessed in four sections 50 μm apart, beginning at the opening of the aortic valves (**F**). Atherosclerotic lesion severity was graded (**G**) and representative images of the haematoxylin-phloxine-saffron stained cross sections of the aortic root are shown (**H**). Data are presented as means ± SEM (n= 14-16), *P<0.05, ***P<0.001 according to one-way ANOVA (E) or two-way ANOVA (F ang G). *E-RUN vs. L-RUN, ^#^SED vs. L-RUN. E-SED data points are labeled white and L-SED data points dark grey within the SED group.

### 3.3 Exercise training does not alter vascular inflammation but influences levels of circulating cytokines

To identify the underlying cause of the reduction of atherosclerotic lesions with late exercise training we assessed the potential modulation of vascular inflammation. A macrophage staining in type III lesions in the aortic root showed a strong trend towards reduced abundance of macrophages in the lesions of L-RUN compared to E-RUN (Fig. 3A and B). However, relative to the total lesion area no difference in macrophage area was observed between the groups. Gene expression analysis of markers involved in leukocyte recruitment (*Ccr2, Icam-1, Vcam-1*), the oxidative stress response (*Gpx1, Sod1*), hypoxia (*Hif1α*) or macrophage abundance and activation (*Adgre1, Cd163*) in the aortic arch did not show differences between groups (Fig. 3C). As not to mask diurnal oscillations in the expression of these genes in tissues collected either at ZT13 (E-SED and E-RUN) or ZT22 (L-SED and L-RUN), the SED groups are shown separately. Circulating levels of the pro-inflammatory cytokines IL-1β and TNFα were not different between groups or timepoints (Fig. 3D, E) while the levels of IL-6 were overall higher at ZT22 than at ZT13 regardless of exercise (Fig. 3F). As the blood samples were collected 24 hours after the last training bout of the exercising mice, no acute spike of skeletal muscle-derived IL-6 was to be expected. Interestingly, an increase in plasma levels of the anti-inflammatory cytokine IL-10 as well as the pro-inflammatory endothelial adhesion molecules ICAM-1 and VCAM-1 observed in L-SED at the end of the active phase at ZT22 as compared to E-SED at ZT13 was counteracted by late exercise training (Fig. 3G-I).

**Figure 3.**
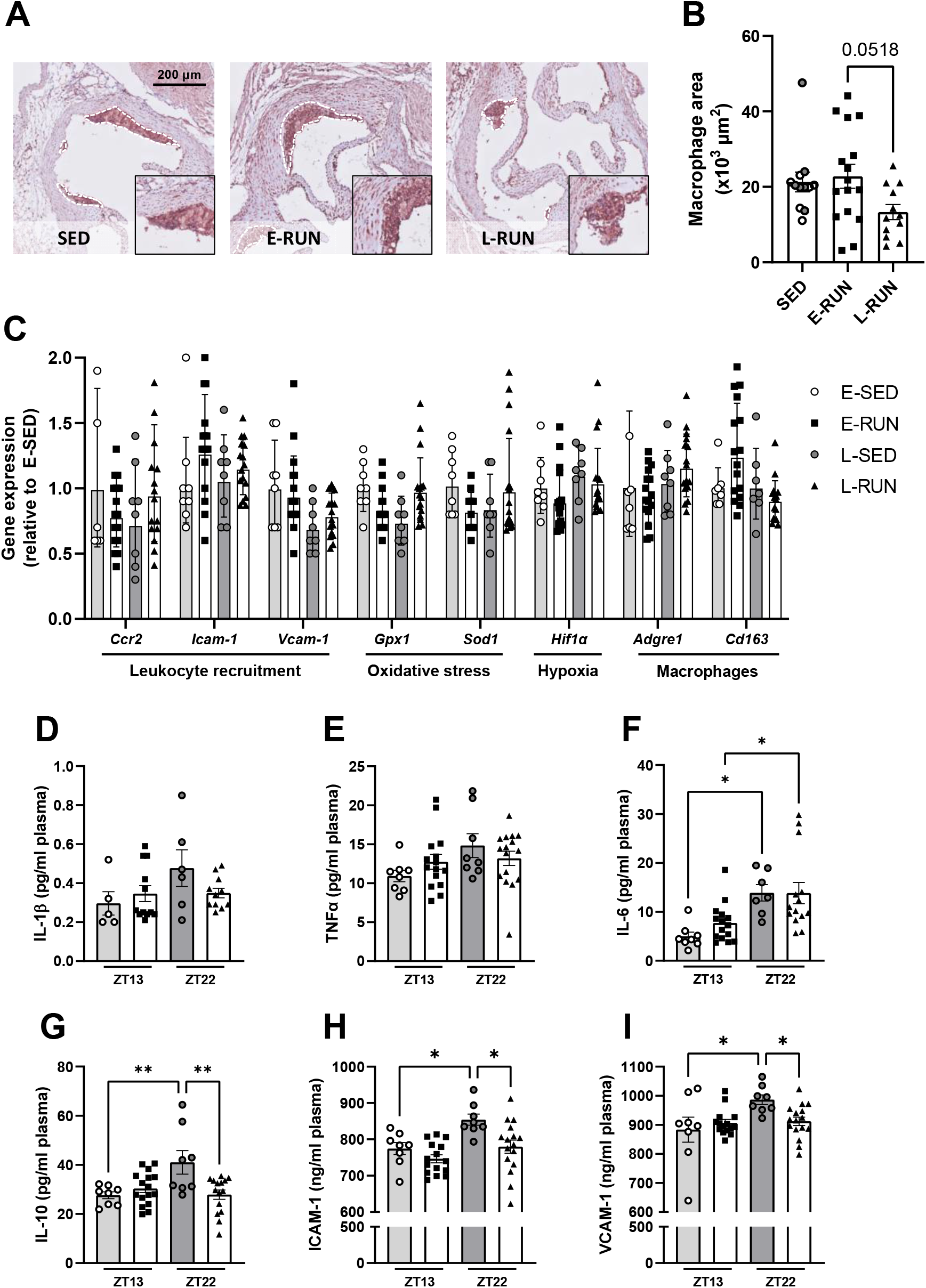
Exercise training does not alter vascular inflammation but influences levels of circulating cytokines. Macrophages in the atherosclerotic lesions in the aortic root were stained with a Mac3 antibody and representative images are shown (**A**). The macrophage area was quantified (**B**) and the gene expression of inflammatory modulators was measured in the aortic tissue (**C**). Plasma levels of the cytokines IL-1β (**D**), TNFα (**E**), IL-6 (**F**), IL-10 (**G**), ICAM-1 (H) and VCAM-1 were assessed in blood collected 24 hours after the last treadmill training bout. Gene expression data (C) is shown as means ± SD (n=8-16), other data are presented as means ± SEM (n= 5-16), *P<0.05, **P<0.01 according to one-way ANOVA. E-SED data points are labeled white and L-SED data points dark grey within the SED group of B.

### 3.4 Circadian timing of handling stress and exercise training affects activity levels of mice and modulates lipid fluxes

For detailed insights into the effect of timed exercise on voluntary locomotor activity, food intake and energy expenditure we performed a follow-up experiment with the same study set-up where the mice were housed in metabolic cages during the fourth week of training. The timing of exercise but surprisingly also handling and temporarily transferring mice to empty cages (as experienced by SED) had an impact on daily behavior. Both E-RUN and E-SED showed reduced locomotor activity during the dark phase after being returned to their home cages compared to L-RUN and L-SED (Fig. 4A). Both exercising groups ate significantly more than their sedentary counterparts but the daily food intake of L-RUN was similar to that of E-RUN (Fig. 4B). Energy expenditure as well as the respiratory exchange ratio (RER) were unchanged between exercising and sedentary mice during the light and dark period and differences between day and night were observed as expected (Fig. 4C, D). Body weight and body composition of these animals along with the real time curves of all parameters measured in the metabolic cages (not normalized as well as normalized to lean body mass) over four days can be found in Supplementary Figures S2-4. Plasma glucose levels were unchanged between training and sedentary animals directly after training as well as after 4, 9 and 13 hours (Supplementary Fig. S2D).

**Figure 4.**
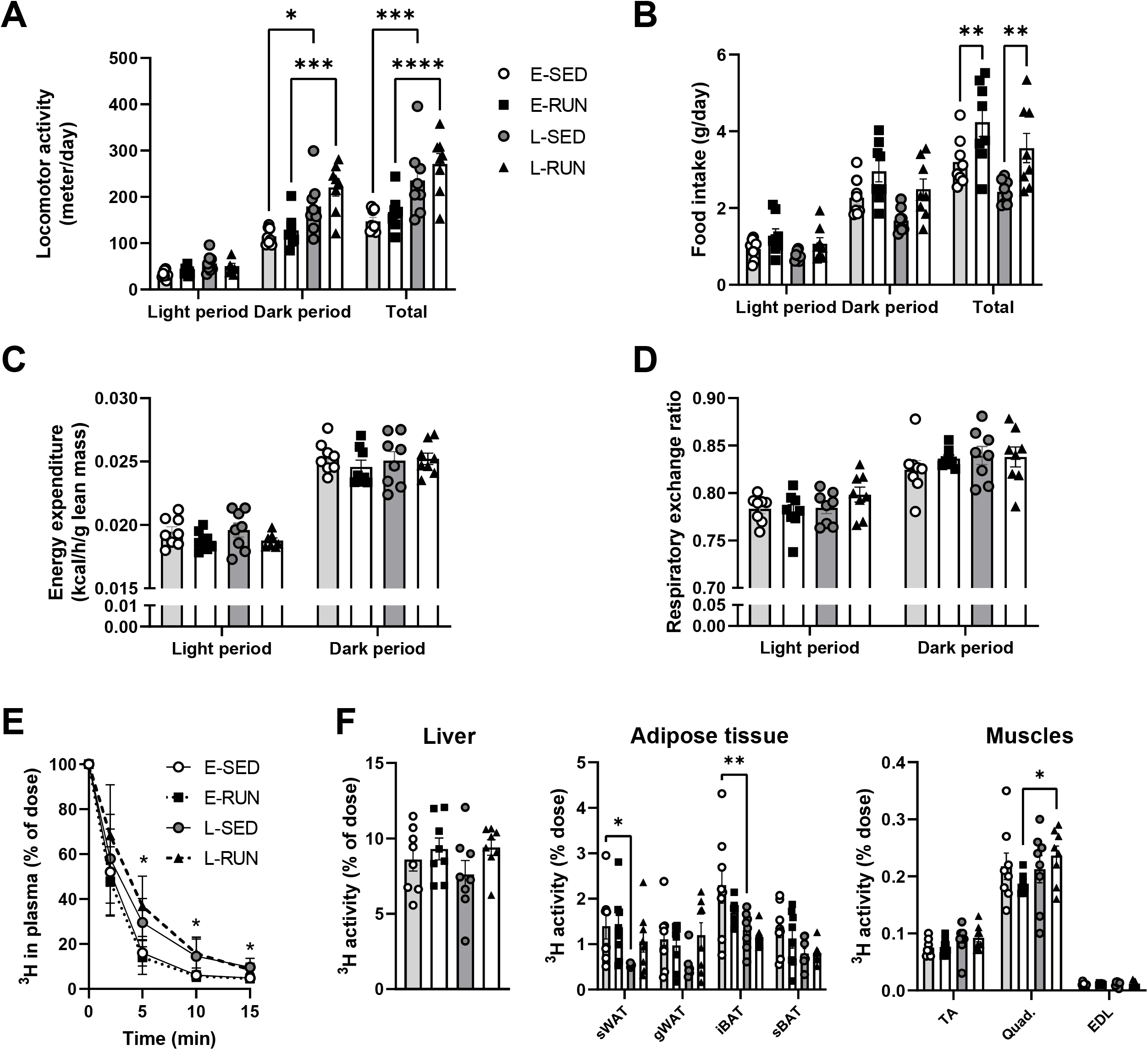
Circadian timing of handling stress and exercise training affects food intake and modulates lipid fluxes. In a follow-up study with the same exercise protocol, APOE*3-Leiden.CETP mice on a cholesterol-rich Western-type diet were housed in metabolic cages during the fourth week of training. Voluntary locomotor activity (**A**) and food intake (**B**) were continuously measured and energy expenditure (**C**) and the respiratory exchange ratio (**D**) were calculated based on O_2_ consumption and CO_2_ production. After six weeks, mice were injected with VLDL-like TG-rich particles labeled with glycerol tri[^3^H]oleate immediately after the last treadmill running bout at ZT14 and ZT23, respectively, and plasma clearance (**E**) and tissue uptake of glycerol tri[^3^H]oleate was assessed (**F**). Data are presented as means ± SEM (n= 6-8), *P<0.05, **P<0.01, ****P<0.0001 according to two-way ANOVA. In E, *E-RUN vs. L-RUN.

Two weeks later, after six weeks of training, we injected the mice in the follow-up study with TG-rich VLDL-like particles labeled with glycerol tri[^3^H]oleate to assess the plasma clearance and organ distribution of TG-derived fatty acids directly after the last exercise bout at ZT14 and ZT23, respectively. At these timepoints, L-RUN had significantly higher plasma TG levels following the exercise bout than E-RUN (E-SED 2.04 ± 0.24, E-RUN 3.04 ± 0.37, L-SED 3.53 ± 0.53, L-RUN 5.16 ± 0.52 mM). No difference was observed in glycerol tri[^3^H]oleate clearance between sedentary and exercising mice but overall, E-RUN cleared lipids from the blood faster than L-RUN and a similar trend was seen for E-SED compared to L-SED (Fig. 4E). Hepatic uptake of [^3^H]oleate was not different between groups but uptake by subcutaneous white adipose tissue (sWAT) and interscapular brown adipose tissue (iBAT) was higher in E-SED than in L-SED while no differences were observed between the exercising groups. In quadriceps muscle, however, the uptake by L-RUN was increased compared to that of E-RUN, albeit not higher than in L-SED (Fig. 4F).

### 3.5 Late exercise training promotes the enrichment of short-chain fatty acid producing gut bacteria

Regular exercise training promotes a change of the composition of the gut microbiota and the microbiota-host-interaction has been shown to modulate the innate immune system that is involved in the development of CVD [23-25]. As the abundance as well as the metabolic activity of gut bacteria and immune cells throughout the body also underly circadian fluctuations it is conceivable that the timing of exercise has a differential impact on the development of inflammatory diseases like atherosclerosis. Accordingly, we investigated h[25ow timed exercise modulates the composition of the gut microbiota in the cecum content with 16S sequencing as well as metagenomic sequencing. Surprisingly, we found that late exercise significantly decreased the alpha diversity, i.e. the diversity of bacteria in the gut of individual mice, when compared to E-RUN and SED (Fig. 5A, 16S sequencing, E-SED and L-SED are shown combined due to overlapping clusters). However, late, but not early, exercise promoted the enrichment of *Firmicutes* (Fig. 5B, 16S sequencing) and particularly the families *Ruminococcaceae* and *Lachnospiraceae* (Fig. 5C, 16S sequencing). These findings were further validated with shotgun metagenomic sequencing of a subset of samples and a principal component analysis confirmed the separate clustering of L-RUN from SED and E-RUN (Fig. 5D). *Firmicutes* were confirmed to be enriched by late training only (Fig. 5E). Species analysis revealed one uncultured *Ruminococcus* species, one uncultured *Lachnospiricea* species, *Lachnospiracea bacterium 28-4*, one uncultured *Clostridiales* species and *Anaerotruncus G3* to be enriched with late exercise in comparison to E-RUN or SED. We furthermore observed a specific enrichment of carbohydrate-active enzymes in the metagenomic dataset of L-RUN which is in line with the enrichment of known fiber-processing bacterial geni involved in short chain fatty acid (SCFA) production (Fig. 5G, H). While we were unable to quantify SCFA plasma levels, we observed a matching increase of the gene expression of the SCFA receptor Olfr78 in oxidative soleus muscle and a similar tendency in the mixed tibialis anterior muscle (TA) in L-RUN compared to E-RUN (p=0.1) (Fig. 5I). A linear regression analysis predicting the modulatory impact of cecal *Ruminococcacea* abundance on atherosclerotic lesion area revealed a significant negative correlation, albeit with weak predictive value (Fig. 5J).

**Figure 5.**
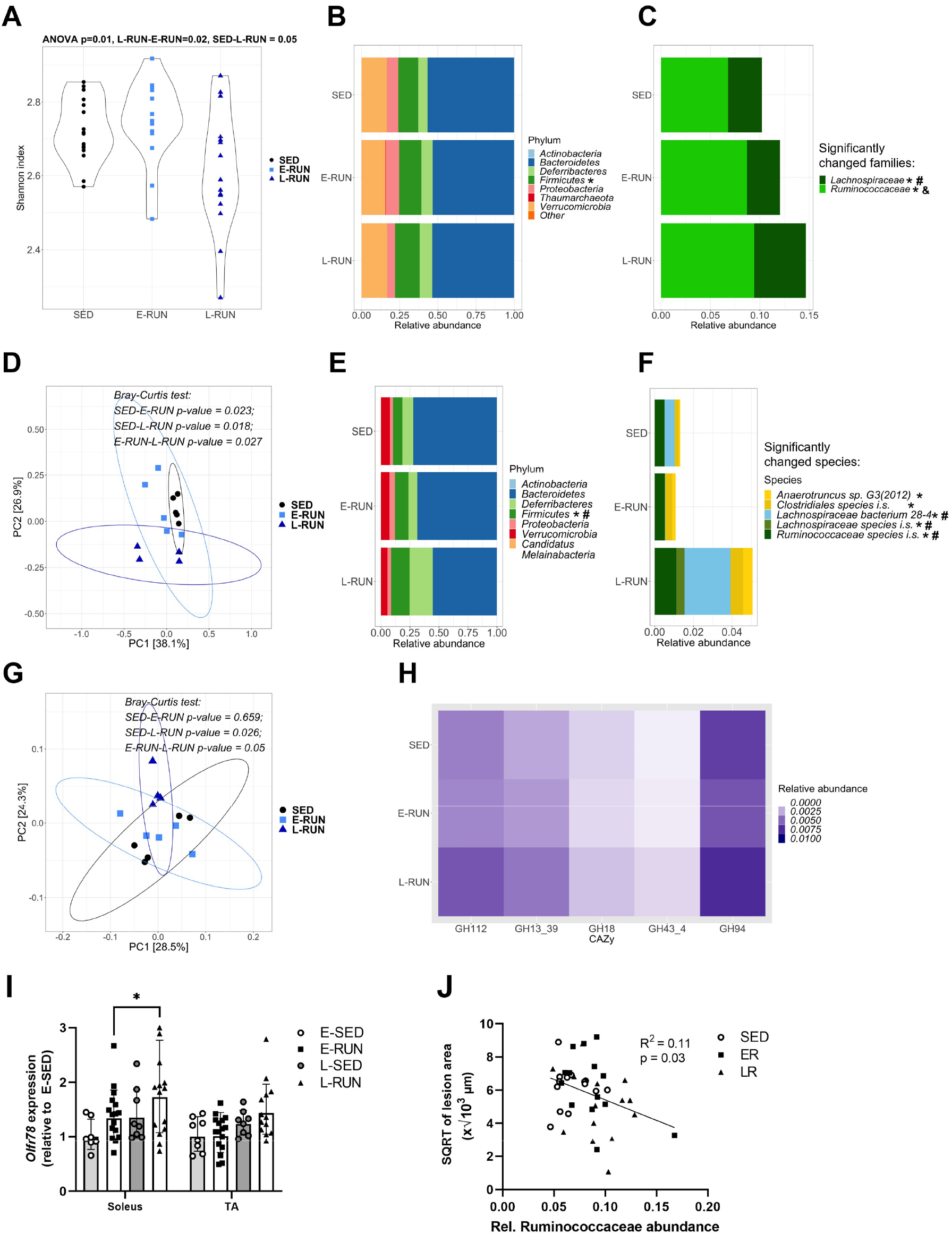
Late exercise training promotes the enrichment of short-chain fatty acid producing gut bacteria. After four weeks of training, the composition of the microbiota in the cecum was analyzed. Based on 16S sequencing, the Shannon diversity index was calculated (**A**) and the bacterial phylae (**B**) and significantly changed families are shown (**C**). Metagenomic shotgun sequencing was performed and a principal component analysis confirmed separate clusters of the groups (**D**). Bacterial phylae (**E**) and significantly changed species are shown (**F**). The abundance of genes encoding carbohydrate-active enzymes (CAZy) was assessed (**G**) and the enriched pathways are shown (**H**). The gene expression of the short-chain fatty acid receptor Olfr78 was quantified in skeletal muscle (**I**) and a linear regression analysis was performed to assess whether atherosclerotic lesion size is a variable of abundance of *Ruminococcacea* (**J**). Gene expression data (**I**) is shown as means ± SD (n=7-16); n=15-16 for 16S sequencing (A-C); n=4-5 for metagenomics sequencing. *P<0.05 according to one-way ANOVA. *SED vs. L-RUN, ^#^E-RUN vs. L-RUN, ^&^SED vs. E-RUN.

## 4. Discussion

All pathophysiological factors contributing to the development of atherosclerosis are under the control of the circadian clock and therefore presumably modifiable through external circadian cues such as physical activity. Exercise training is well known to improve cardiometabolic health, but it has not been elucidated yet whether the timing of exercise training affects its beneficial cardiovascular outcomes. Here we demonstrate for the first time that exercise training at different times during the active phase of mice has a differential impact on the development of atherosclerosis.

Late exercise training ameliorated the gain of body weight throughout four weeks of treadmill running compared to early training. This was primarily due to a pronounced loss of fat mass only observed in mice training late. These findings are in line with previous reports showing that voluntary wheel running in the late active phase protects mice against diet-induced obesity while early active phase running does not [26, 27]. As differences in food intake observed here between the two sedentary groups did not translate into differences in fat mass, this may suggest that mice exercising late are less efficient in extracting energy from nutrients. Endurance training in mice generally does not evoke a gain in lean body mass [28]. Instead, muscle mass correlates with the overall body weight of mice, explaining the reduced gain of lean mass in both exercising groups compared to their overall heavier untrained counterparts.

Interestingly, four weeks of treadmill training did not lower plasma lipid levels in these dyslipidemic animals, but late training independently of changes in cholesterol exposure reduced the development of atherosclerotic lesions. While clinical studies often, but not always, report improved plasma TG levels following exercise intervention studies, generally no or minor reductions of plasma cholesterol levels are reported [29, 30]. It is conceivable, however, that in the present study the exercise training was unable to counteract the continuous high influx of dietary lipids from the cholesterol-enriched WTD, or that short transient changes in whole body lipid metabolism occurred during the training and recovery that contributed to the reduced lesion formation. Metabolomics analyses in muscle, liver and adipose tissue in lean healthy mice showed that lipid metabolites are among the most altered metabolite classes following acute exercise and that the magnitude of these metabolite level changes greatly depends on the time of day [10]. Hereby, the nutritional status matters as the body has to activate alternative energy sources such as ketone bodies when intramuscular and hepatic energy stores are depleted coming out of the rest phase. Conversely, during late active phase training energy stores are full and liver and adipose tissue can provide energy for the exercising muscles via glycogen and lipid degradation [9]. When assessing the plasma clearance and tissue uptake of radioactively labeled TG by peripheral tissues directly after the last training bout we indeed observed time-of-day specific differences. TG clearance at the end of the active phase was slower than in the early active phase and subcutaneous white fat and brown fat of sedentary mice took up less TG, presumably due to full energy stores and lower BAT activity at the end of the active phase [31]. At the same time, the TG uptake by quadriceps muscle was increased in late trained mice following exercise as expected as the muscle is supposedly primed to (re)fuel via lipid uptake. While plasma glucose levels were unaffected, we cannot rule out that improved insulin sensitivity with late exercise contributed to the observed reduction of atherosclerotic lesion burden.

The progression of atherosclerosis can be modulated either through the reduction of plasma cholesterol levels or of the inflammatory processes contributing to lesion development. Curiously, when assessing the relative macrophage content of the aortic lesions we found no differences between the groups and also gene expression analyses of markers of leukocyte recruitment and oxidative stress in the aorta provided no evidence of inflammatory improvement with late training. However, we did not directly assess the activation levels of the lesional macrophages which may have differed and contributed to altered lesion progression. Circulating IL-6 levels were not different between sedentary and trained mice 24 h after the last exercise bout but a diurnal effect was observed with higher IL-6 levels at the end of the active phase, matching the rhythm found in humans [32]. IL-6 has both pro- and anti-inflammatory properties depending on its signaling modality and is often considered detrimental in atherosclerosis development [33]. However, it has also been reported that the administration of IL-6 can reduce the atherosclerotic lesion burden [34]. Of note, exercise also increases the circulating levels of sgp130, an endogenous inhibitor of IL-6’s pro-inflammatory signaling, that reduces atherosclerosis development when administered exogenously, but it remains unclear to what extent this effect depends on time-of-day [35, 36]. Here, it can be speculated that an acute exercise-induced increase of IL-6 late in the active phase that coincides with the highest abundance of the membrane-bound IL-6 receptor on immune cells and an increase in circulating sgp130 may be atheroprotective [37, 38]. Unexpectedly, late training prevented a diurnal increase of IL-10, a cytokine that has long been established to be anti-inflammatory and to ameliorate atherosclerosis [39]. At the same time, late exercise prevented the diurnal increase of the endothelial dysfunction markers ICAM-1 and VCAM-1 at the end of dark phase, possibly suggesting that exercise is more effective when the inflammatory tonus is overall higher. The spatial impact of exercise at different times of day on peaks and overall circulating levels of pro- and anti-inflammatory cytokine levels is, however, still unclear. Additionally, the interaction of timed exercise with inflammatory processes in advanced atherosclerosis should be further investigated.

Although energy expenditure was similar in all groups, voluntary locomotor activity levels differed between the early and late groups. Mice that were handled in the beginning of the active phase – sedentary and trained mice alike – moved less within the home cage during the remainder of the day. However, as the atherosclerosis lesion burden between the early and late sedentary animals was comparable these differences in locomotor activity likely did not contribute to the differences in atherosclerosis observed in the early and late exercising mice. Similarly, both exercising groups ate more over 24 hours than sedentary animals but no differences were seen in food intake between early and late exercising mice, making changes in food intake an unlikely contributing factor to differences seen in atherosclerosis development. While these findings underline the importance of considering handling stress in mouse studies aiming to investigate exercise effects, they do not provide an explanation for the findings on atherosclerosis and fat mass loss.

Recent studies in mice and humans have begun to elucidate the involvement of gut microbiome-host interactions regulating host metabolism [40]. Here we demonstrate that late, but not early, training surprisingly decreased the bacterial diversity within the gut but at the same time promoted the enrichment of SCFA-producing bacteria that have been associated with a modulation of atherosclerosis development in mice [41, 42]. The SCFA derived from bacterial fermentation of dietary fiber can inhibit the formation of foam cells and overall reduce inflammation, in part through the enhancement of intestinal barrier function which blocks the flux of bacterial toxins into the blood [43, 44]. Indeed, it was confirmed recently that the exercise-induced enrichment of SCFA-producing bacteria ameliorated vascular inflammation and atherosclerosis development in ApoE knockout mice [45]. While we were unable to reliably quantify circulating levels of SCFA in cardiac puncture blood due to very low concentration or assess the state of the gut barrier, we found that the late training increased the gene expression of the SCFA receptor Olfr78 in the oxidative soleus muscle. This could suggest higher circulating levels of SCFA or a higher peripheral sensitivity towards SCFA with late exercise. Whether the time-of-day specific effect of exercise on the gut microbiome is causal and mechanistically responsible or in part contributing to the observed reduction of atherosclerosis development with late training remains to be elucidated beyond the association reported here. Little is known about the two cultured identified bacterial species upregulated with late training, but *Lachnospiraceae bacterium 28-4* has been linked to the attenuation of obesity in high-fat fed mice [46, 47].

## 5. Conclusion

By using a well-established model of human-like cardiometabolic disease, we show that later exercise training rather than early training attenuates diet-induced gain of fat mass and reduces atherosclerosis development (Fig. 6). Overall, the data presented here underscores that the timing of exercise may be an important parameter in optimizing exercise recommendations for patients with cardiovascular diseases. Future mechanistic studies should identify the role of the gut microbiota in this effect along with further characterizations of the molecular changes within the atherosclerotic lesions.

**Figure 6.**
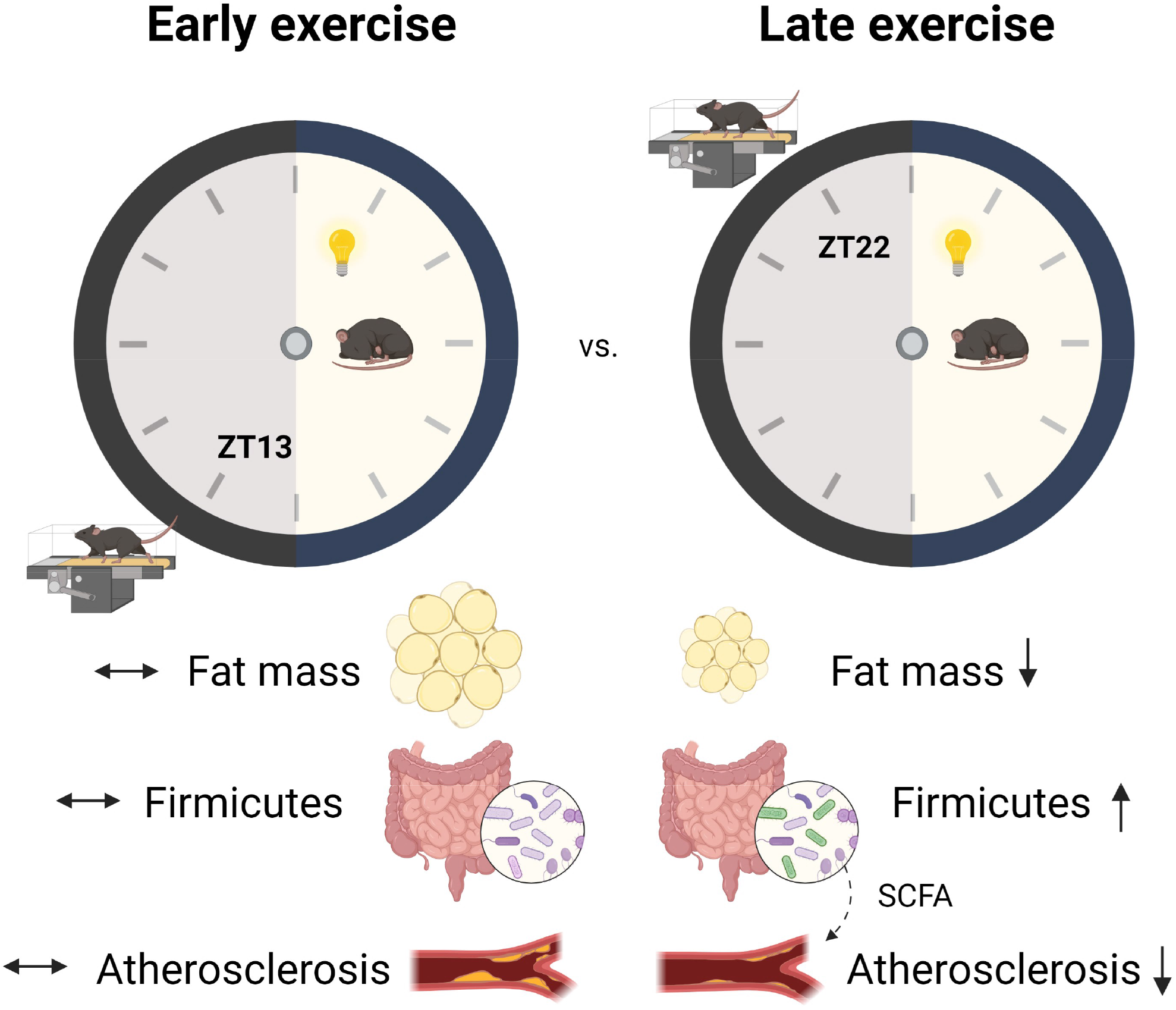
Schematic overview of main findings and proposed mechanism. Arrows indicate experimental findings, the dashed arrow indicates the proposed underlying mechanism. Gut microbiota-derived SCFAs may modulate atherosclerosis directly or via the involvement of other tissues such as skeletal muscle.

## Data availability statement

The 16S sequencing data are available at https://zenodo.org/record/6992206#. Following peer-review the restricted access will be removed. All data is available upon reasonable request to the corresponding author.

## Author contribution

Study conception and design: M.S., Z.Y., P.C.N.R.; data collection: M.S., Z.Y., A.K., W.P., A.B., S.P., A.C.M.P., T.C.M.S.; analysis and interpretation of results: M.S., Z.Y., A.K., M.H., S.K., P.C.N.R.; draft manuscript preparation: M.S., Z.Y., A.K., P.C.N.R. All authors reviewed the results and approved the final version of the manuscript.

## Acknowledgements

We thank Reshma Lalai and Chris van der Bent (Div. of Endocrinology, Dept. of Medicine, LUMC, Leiden, the Netherlands) for their excellent technical assistance. We furthermore thank Aswin Verhoeven and Martin Giera (both Center for Proteomics and Metabolomics, Leiden University Medical Center, Leiden, The Netherlands) for their valuable technical contributions. This study was financed by a grant from the Novo Nordisk Foundation to M.S. (NNF18OC0032394). S.K. was supported by the Dutch Heart Foundation (2017T016). Z.Y. was supported by a full-time PhD scholarship from the China Scholarship Council.

