## Supplemental materials and figures for "Time to run: Late rather than early exercise training in mice remodels the gut microbiome and reduces atherosclerosis development"

### Supplementary methods

#### 16S sequencing

Data processing

The QIIME 2 2021.8 tool [1] was used to process the 16S sequencing data. The raw sequence data were denoised and quality filtered with the DADA2 plugin (via “denoise-paired”) [2]. The obtained representative sequences were classified with the feature-classifier plugin (via “classify-sklearn”) [3] using the pre-trained Silva 138 99% OTUs full-length sequences classifier at 0.99 sequence identity. The resulting raw hits matrix was collected using the “taxa collapse” function.

Statistical analysis

Further statistical analyses were conducted with R version 4.1.1. For the statistical analyses, the Phyloseq package (version 1.36.0) [4] was used to integrate the hits matrix, taxonomy and metadata. To determine alpha-diversity (presented as Shannon index), the package’s function “estimate richness” was used and a one-way ANOVA was performed. To determine beta-diversity, the package’s function “plot_ordination” was used with additional Bray-Curtis testing.

For the identification of specific geni and species increased in one of the groups, DESeq2 (version 1.32, method = “poscounts”, prevalence filtering = 0.25) [5] for raw abundance and SIAMCAT (version 1.12, abundance filtering = 0.01) [6] for relative abundance were used. Results are shown if the adjusted p-value < 0.05 and hits were registered by DESeq2. Ggplot2 (version 3.3.5) [7] was used for all visualizations.

#### Metagenomics sequencing

Quality control

The Ngless (version 1.0) [8] framework was used for the pre-processing of the raw metagenomics reads derived from shotgun sequencing. Reads containing less than 45 nucleotides with a quality score above 25 were discarded. Then, reads were mapped against the mouse genome database “mocat” (version 0.0, module of Ngless, reference = “mm10.p5") and reads with more than 90% similarity were also discarded.

Abundance matrix generation

A mOTUs species matrix was generated using the mOTU-tool (version 2.6) [9] , a built-in module of Ngless. Further statistical analyses were conducted with R version 4.1.1. For the statistical analyses, the Phyloseq package (version 1.36.0) [4] was used to integrate the hits matrix, taxonomy and metadata. To determine alpha-diversity (presented as Shannon index), the package’s function “estimate richness” was used and a one-way ANOVA was performed. To determine beta-diversity, the package’s function “plot_ordination” was used with additional Bray-Curtis testing.

For the identification of species and mOTUs increased in one of the groups, DESeq2 (version 1.32, method = “poscounts”, prevalence filtering = 0.25) [5] for raw abundances and Corncob (version 0.2.0, abundance filtering = 0.01) [10] for relative abundances were used. Results are shown if adjusted p-value < 0.05. Ggplot2 (version 3.3.5) [7] was used for all visualizations.

CAZy identification

For the mapping of carbohydrate-active enzymes (CAZy), the dbCAN2 tool [11] was used. As the tool requires long contigs instead of short sequences, contigs were assembled using the IDBA-UD tool. "Contig.fa" output was used as the most optimal contig outcome of the run. Dbcan output consists of three hit matrixes – GMMER, Hotpep and DIAMOND. To ensure that only true hits are extracted, a contig was considered a hit only if it was detected by all three databases. The resulting CAZy abundance matrix was analyzed using LEFse to find CAZy families increased in any of the groups.

#### Supplementary Figure legends

**Supplementary Figure 1: Chronic stress exposure.** After four weeks of training, the adrenal glands were weighed (A). Data are presented as means ± SEM (n= 8-16). E-SED data points are labeled white and L-SED data points dark grey within the SED group.

**Supplementary Figure 2: Body composition and plasma glucose levels.** After five weeks of training, in a second mouse cohort, body weight (A), lean mass (B) and fat mass (C) was assessed before the mice were housed in metabolic cages. Two weeks prior, during week 3 of the training program, plasma glucose levels were measured over 13 hours following a training bout (D). The shaded area indicates the dark phase and arrows indicate the training timepoints (E-RUN, white, ZT 13; L-RUN, dark grey, ZT22). Data are presented as means ± SEM (n= 8).

**Supplementary Figure 3:** After five weeks of training, animals of a second cohort were housed in metabolic cages and oxygen consumption, carbon dioxide production and energy expenditure over four days are shown not normalized (left) and normalized to lean mass (right). Shaded areas indicate the dark phases. Realtime data are presented as group means (n= 8).

**Supplementary Figure 4:** After five weeks of training, animals of a second cohort were housed in metabolic cages and fat and carbohydrate oxidation rates not normalized (left) and normalized to lean mass (right) as well as the respiratory exchange ratio and locomotor activity over four days are shown. Shaded areas indicate the dark phases. Realtime data are presented as group means (n= 8).

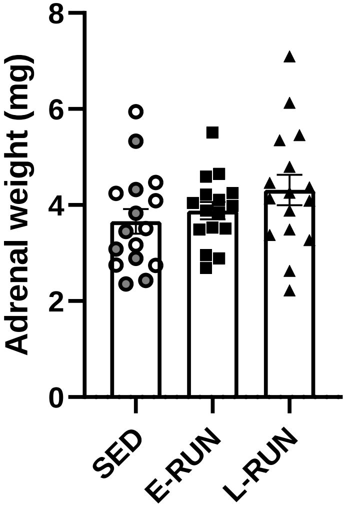
Adrenal weight (mg)
0
2
4
6
8
SED
E-RUN
L-RUN

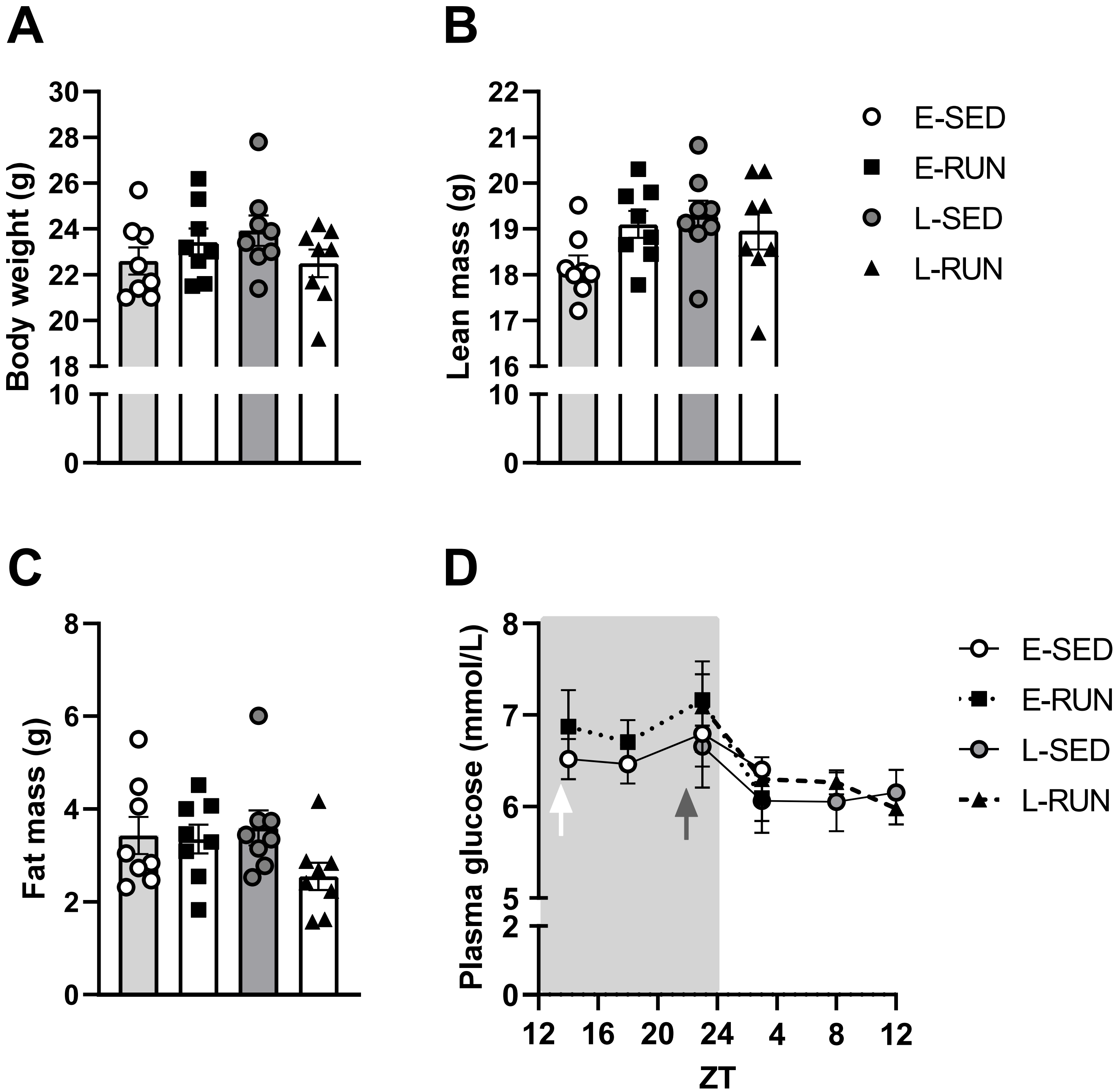
A
B
C
D
Body weight (g)
Lean mass (g)
Fat mass (g)
Plasma glucose (mmol/L)
ZT
E-SED
E-RUN
L-SED
L-RUN

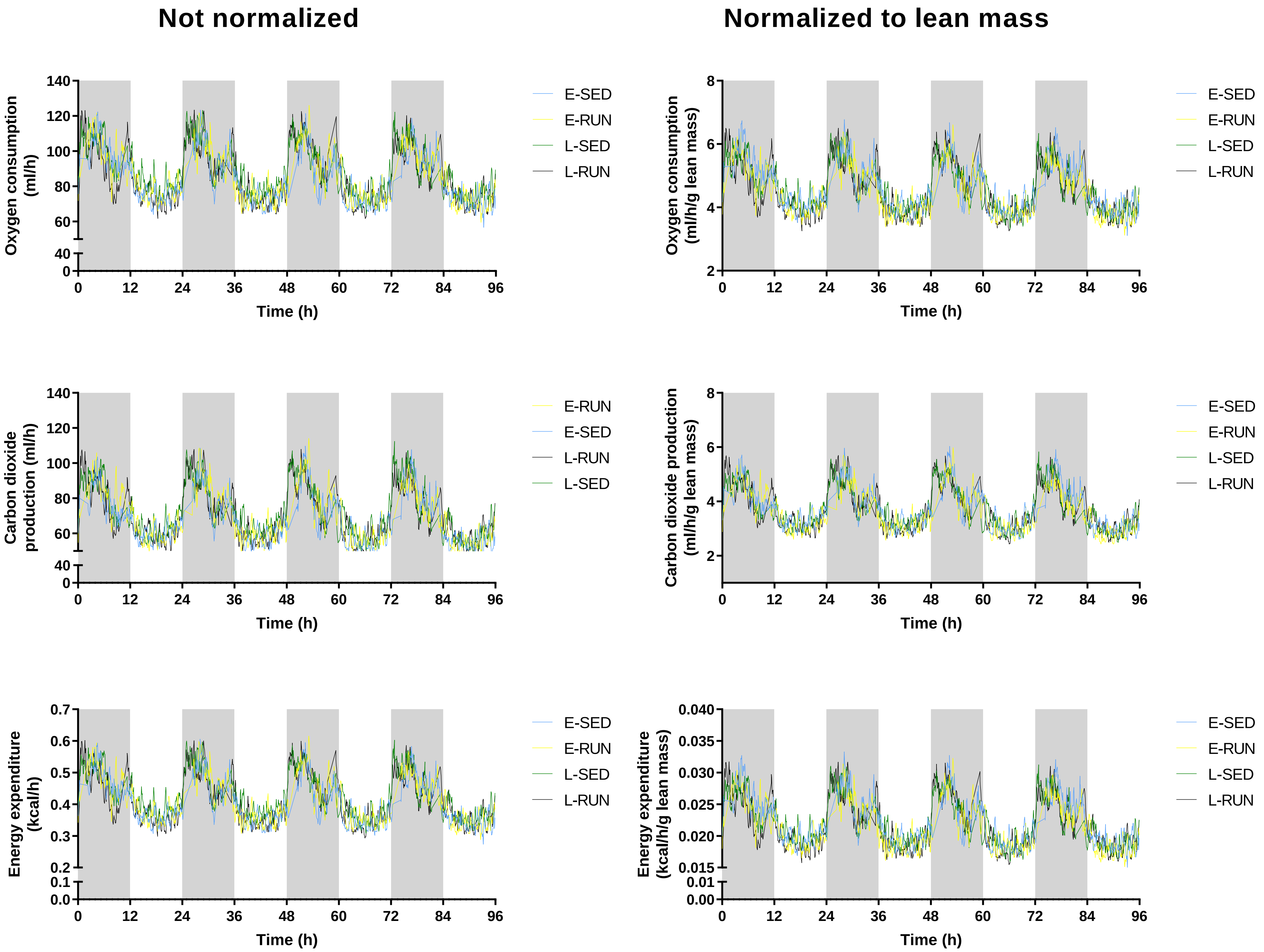
Not normalized
Normalized to lean mass
Oxygen consumption (ml/h)
Oxygen consumption (ml/h/g lean mass)
Carbon dioxide production (ml/h)
Carbon dioxide production (ml/h/g lean mass)
Energy expenditure (kcal/h)
Energy expenditure (kcal/h/g lean mass)
Time (h)
0 12 24 36 48 60 72 84 96
0 40 60 80 100 120 140
2 4 6 8
0.0 0.1 0.2 0.3 0.4 0.5 0.6 0.7
0.00 0.01 0.015 0.020 0.025 0.030 0.035 0.040
E-SED
E-RUN
L-SED
L-RUN

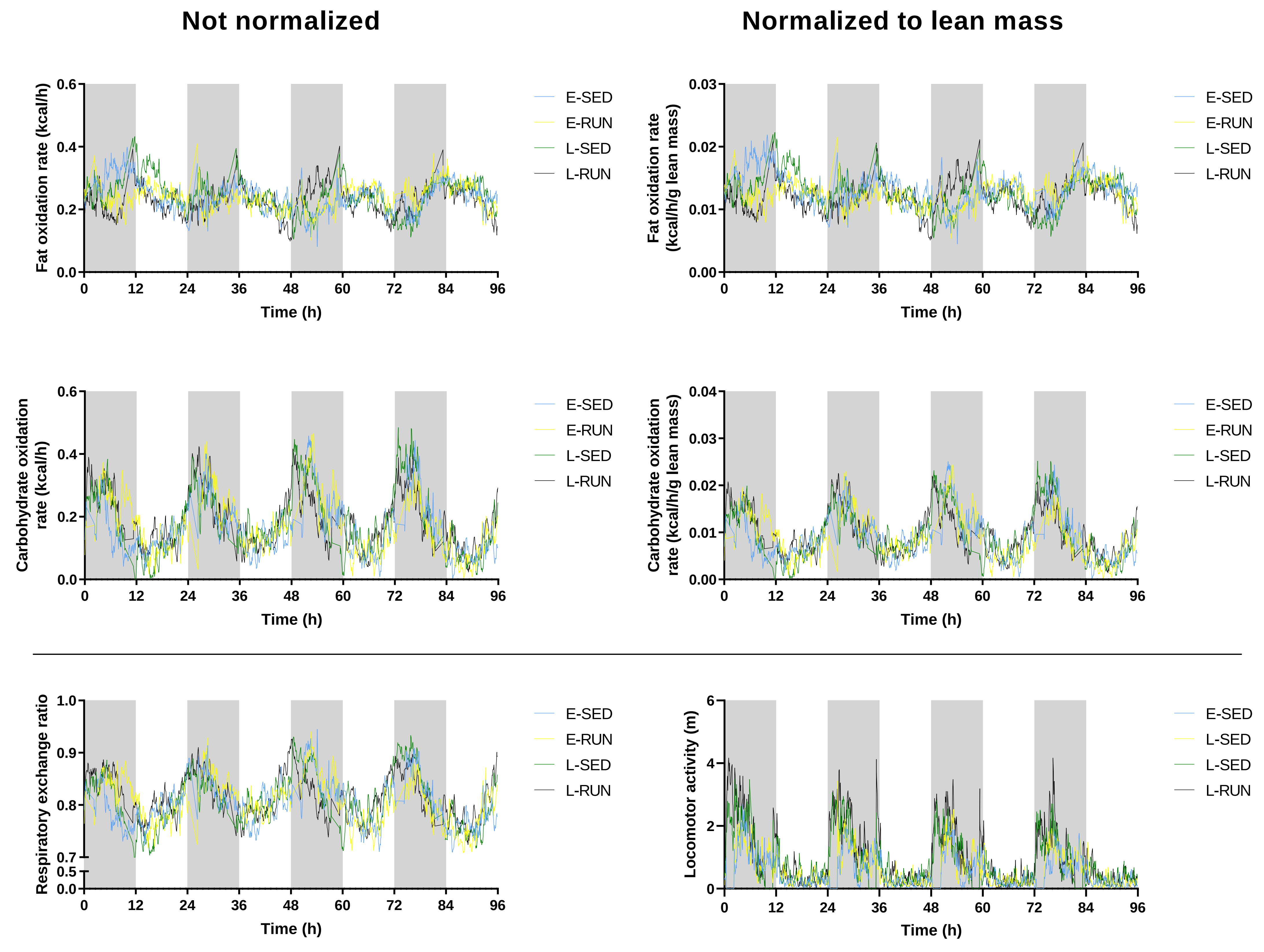
Not normalized
Normalized to lean mass
Fat oxidation rate (kcal/h)
Fat oxidation rate (kcal/h/g lean mass)
Carbohydrate oxidation rate (kcal/h)
Carbohydrate oxidation rate (kcal/h/g lean mass)
Respiratory exchange ratio
Locomotor activity (m)
Time (h)
0 12 24 36 48 60 72 84 96
E-SED
E-RUN
L-SED
L-RUN
E-SED
L-SED
L-SED
L-RUN
